# A de novo approach to inferring within-host fitness effects during untreated HIV-1 infection

**DOI:** 10.1101/825117

**Authors:** Christopher J. R. Illingworth, Jayna Raghwani, David Serwadda, Nelson K. Sewankambo, Merlin L. Robb, Michael A. Eller, Andrew R. Redd, Thomas C. Quinn, Katrina A. Lythgoe

## Abstract

In the absence of effective antiviral therapy, HIV-1 evolves in response to the within-host environment, of which the immune system is an important aspect. During the earliest stages of infection, this process of evolution is very rapid, driven by a small number of CTL escape mechanisms. As the infection progresses, immune escape variants evolve under reduced magnitudes of selection, while competition between an increasing number of polymorphic alleles (i.e., clonal interference) makes it difficult to quantify the magnitude of selection acting upon specific variant alleles. To tackle this complex problem, we developed a novel multi-locus inference method to evaluate the role of selection during the chronic stage of within-host infection. We applied this method to targeted sequence data from the p24 and gp41 regions of HIV-1 collected from 34 patients with long-term untreated HIV-1 infection. We identify a broad distribution of beneficial fitness effects during infection, with a small number of variants evolving under strong selection and very many variants evolving under weaker selection. The uniquely large number of infections analysed granted a previously unparalleled statistical power to identify loci at which selection could be inferred to act with statistical confidence. Our model makes no prior assumptions about the nature of alleles under selection, such that any synonymous or non-synonymous variant may be inferred to evolve under selection. However, the majority of variants inferred with confidence to be under selection were non-synonymous in nature, and in nearly all cases were associated with either CTL escape in p24 or neutralising antibody escape in gp41. Sites inferred to be under selection in multiple hosts have high within-host and between-host diversity albeit not all sites with high between-host diversity were inferred to be under selection at the within-host level. Our identification of selection at sites associated with resistance to broadly neutralising antibodies (bNAbs) highlights the need to fully understand the role of selection in untreated individuals when designing bNAb based therapies.

**Author Summary:** During the within-host evolution of HIV-1, the diversity of the viral population increases, with many beneficial variants competing against each other. This competition, known as clonal interference, makes the identification of variants under positive selection a challenging task. We here apply a novel method for the inference of selection to targeted within-host sequence data describing changes in the p24 and gp41 genes during HIV-1 infection in 34 patients. Our method adopts a parsimonious approach, assigning selection to the smallest number of variants necessary to explain the evolution of the system. The large size of our dataset allows for the confident identification of variants under selection, alleles at certain loci being repeatedly inferred as under selection within multiple individuals. While early CTL escape mutations have been identified to evolve under strong positive selection, we identify a distribution of beneficial fitness effects in which a large number of mutations are under weak selection. Variants that were confidently identified under selection were primarily found to be associated with either CTL escape in p24 or neutralising antibody escape in gp41, including sites associated with escape from broadly neutralising antibodies. We find that the most frequently selected loci have high diversity both within-host and at the between-host level.

## Introduction

In the absence of effective antiretroviral therapy, HIV-1 evolves rapidly during infection. A key driver of evolution is the influence of the host immune system; cytotoxic CD8+ T-cells (CTLs) and neutralising antibodies (nAbs) impose selection on the virus, leading to the emergence of immune escape mutations[1]. However, other factors also influence viral evolution. For example, the host-specific nature of the immune response leads to the accumulation of mutations which are deleterious to the virus upon transmission to a new host. During the course of a new infection such variants are often lost, in particular where they occur at sites which in general are under strong purifying selection[2,3]. Selection may further act for protein or RNA secondary structure[4,5].

The complex nature of selection has led to multiple studies evaluating how the viral genotype may be both constrained and shaped during the course of evolution. These include the use of techniques for *in vitro* mutagenesis, and analyses of viral sequence data, evaluated at the level of population consensus or through deep sequencing exploring within-host variation at one or more time points during infection. For example, mutagenesis of HIV-1 proteins has allowed the measurement *in vitro* of the effect of specific mutations [6]. The development of technologies for high-throughput mutagenesis has enabled such measurements to be made across very large sets of potential mutations[7–9]. Mathematical methods combining such results have been used to generate an overview of fitness costs and epistatic effects for the virus[10]. Measurements of this form provide a base-level estimation of the general fitness landscape of the virus, albeit that *in vitro* data may be limited in the extent to which it captures the behaviour of the virus within a human host.

Many years of study of HIV-1 have led to the collection of consensus genome sequence data for a large number of individual infections[11]. Such data have allowed techniques such as the fitting of maximum entropy models, which characterise the fitness costs of non-consensus variants in regions of the viral genome[12–14]. In this case, the fitness effects learned give a picture of the generic landscape upon which HIV-1 evolves, averaged over all within-host environments represented by the sequence database, albeit those host-specific environments may differ[15,16]. The extent of conservation at a particular genetic locus indicates the extent to which purifying selection acts upon the majority allele[17].

Short-read deep-sequencing data has provided valuable insights into how fitness effects shape the evolution of HIV-1. Studies can be broadly categorised into those that consider purifying selection, and those that consider positive selection. Purifying (or negative) selection represents the process by which deleterious variants are purged from a population. Over time the frequency of a variant under purifying selection evolves in a statistically predictable way towards an equilibrium state via mutation-selection balance[18]. Exploiting this fact, changes in allele frequency observed over time from across the course of untreated infections[2], or comparing single time points across multiple individuals, have been used to estimate the magnitude of selection and the mutation rate acting upon distinct regions of the genome[19][17].

Positive selection represents the process by which favourable variants are driven towards fixation, and as with purifying selection population genetic methods can be adopted for the inference of fitness effects. For example, a series of models have been developed for the inference of HIV-1 escape rates from CTL responses. Whereas earlier approaches to this problem considered viral escape from a single CTL response[20–23], more recent studies have considered the multiple immune responses that arise successively during infection [24–26]. Under such circumstances, interference between beneficial viral mutations affects the population dynamics [27]. Therefore, accounting for this clonal interference is critical if the role of selection is to be correctly inferred[28–30].

Studies assessing fitness effects in within-host HIV-1 infection have often focused upon the earliest stages of infection when strong selection on CTL escape mutations typically dominates the viral population dynamics[24,30]; in this circumstance, we can model evolution as a competition between a relatively small number of viral genotypes [24,31]. Later in infection, where escape mutations are less strongly beneficial, and where synonymous diversity has had longer to accumulate [2], the potential for hitchhiking and clonal interference is greater, such that variants observed at high frequency are less certain to have evolved under positive selection. Consequently, distinguishing selected from non-selected variants poses a substantial challenge. To address this, we present a *de novo* approach for inferring selection from HIV-1 sequence data in which any variant allele may, in theory, be detected as under selection. We adopt a parsimonious approach, assigning selection to the smallest set of variants required to explain the observed multi-locus sequence data under a likelihood model. Applied to targeted sequence data from a substantial cohort of 34 untreated individuals living in Uganda[32], we determine how selection drives viral evolution. In the presence of pervasive interference between alleles in linkage disequilibrium with one another, our consideration of data from a large number of individuals is fundamental in providing statistical confidence in the assignment of selection. Specifically, the repeated inference of selection at the same locus in independent cases of infection enhances the power of our study to elucidate how selection during individual infections shapes genetic diversity at the population level.

## Results

We applied an evolutionary inference method to deep-sequencing data spanning multiple years of infection from 34 untreated individuals living in Rakai, Uganda, enabling us to infer positive selection acting on part of the gp41 region of *env* (324 base pairs) and the p24 region of *gag* (387 base pairs)[32].

### Extent of selection

An initial application of our method found extensive evidence of positive selection in the viral genome, with 74 (out of 387) nucleotide sites in p24, and 81 (out of 324) nucleotide sites in gp41, being inferred to evolve under positive selection in at least one individual (S1 Table and S1 Fig). Our method explicitly accounts for linkage disequilibrium between alleles observed in the sequencing data[28,33]; however a potential exists for interactions between observed alleles, in the targeted sequence region, and non-observed alleles in flanking regions of the genome. Simulation studies showed that our approach performs well in multi-locus systems where all selected alleles are observed, but is prone to generating false positive inferences of selection where non-observed selected alleles are in linkage disequilibrium with observed alleles (S1 Text). Despite not always attributing selection to the correct locus, the distribution of the inferred strengths of selection, in cases where these could be inferred with confidence, was not statistically distinguishable from the ‘true’ distribution of strengths used to conduct the simulations (S2 Fig; p=0.078, Kolmogorov-Smirnov test). Interpreted at face value, these results reflect the extent of selection in and around the regions of the genome under analysis without necessarily correctly attributing selection to the correct genetic variant.

### Strength and time of onset of selection

Inferred variants generally evolved under weak selection. Taking our inferences at face value, we identified selected variants for which the magnitude of selection could be quantified with some degree of precision. The uncertainty in our estimates depends upon the extent to which data are available; for example, when a variant emerges and fixes between two time points, only a lower bound on the strength of selection can be inferred[34]. For this analysis, we generated confidence intervals for each inferred magnitude of selection using a likelihood-based method, retaining only variants for which the upper and lower bounds of this interval differed by no more than an order of magnitude. Our model assumed that a selected variant evolved under neutrality until some fixed point in time, following which it conferred a constant fitness advantage or disadvantage to the virus. Distributions of fitness effects compiled from the inference showed that most of the identified alleles under selection experienced very weak selective effects, with long tails of alleles evolving under strong positive selection (Fig 1). In p24, around 65% of inferred selected variants had a strength of selection of less than 5% per generation, while in gp41 approximately half of variants were under this threshold. In our model, selection of 5% per generation would cause, in the absence of interference effects, a change in allele frequency from 5% to 95% in a period of just under eight months. Our approach likely underestimates the proportion of variants under weak selection; very weak selection would not produce a change in the population sufficient to be identified from the data. Variants which fixed in the population before the first sample was collected also cannot be identified from our data, such that the extreme beneficial end of the distribution may also be under-represented. Our result, of a preponderance of weakly beneficial mutations during chronic infection, is robust to these two qualifications. Most selection acting upon variants was inferred to kick in during the first year of infection and almost all within two years; we note that the finite length of sequencing could restrict the inference of more lately selected variation.

**Fig 1:**
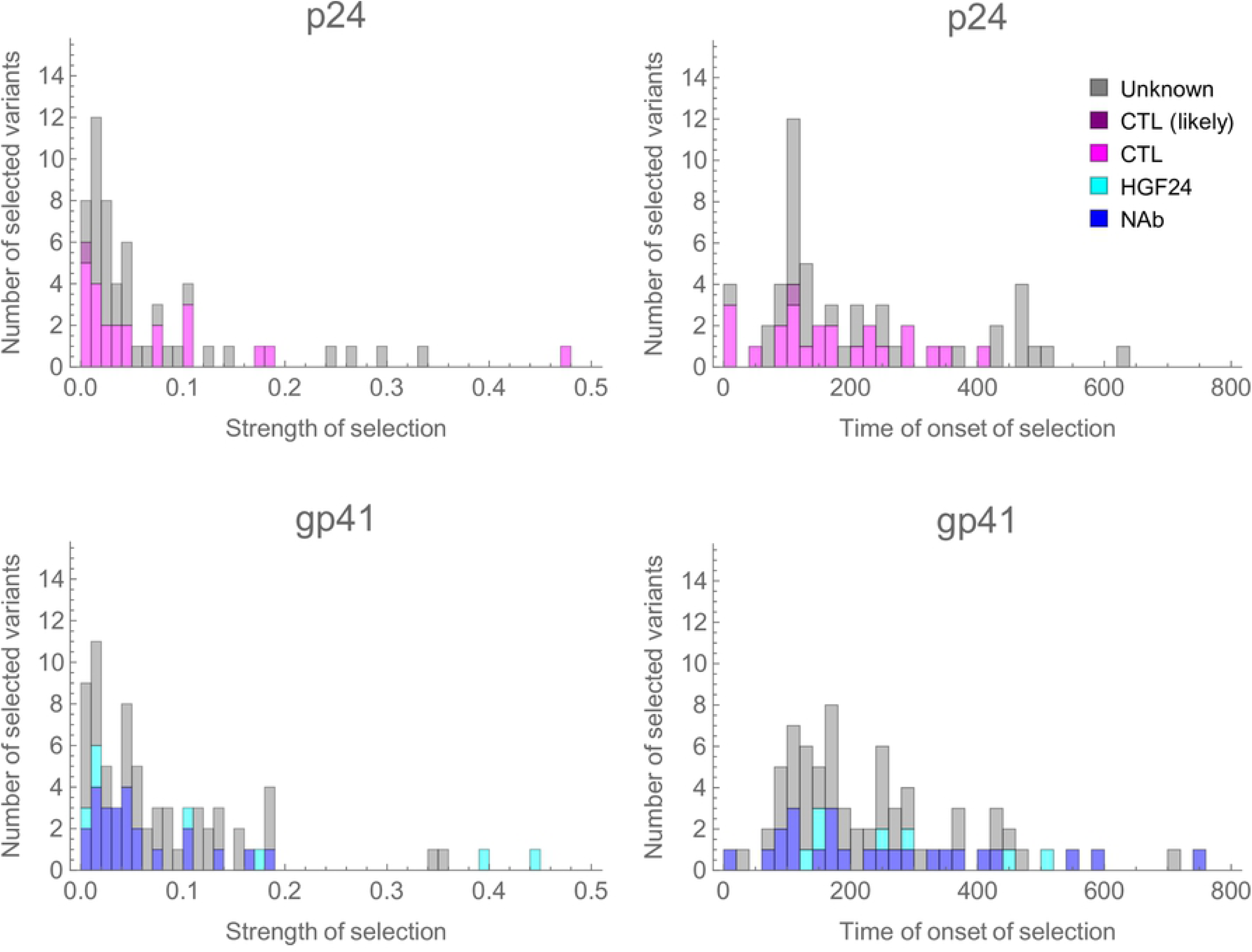
Histograms of inferred strengths of selection and times in days of onset of selection across 34 individuals. The maximum likelihood estimates are shown in each case, for the subset of the data where the upper and lower bound confidence intervals on the strength of selection are within an order of magnitude of each other. Coloured bars indicate variants at nucleotide sites we are confident are under selection in at least one individual and are coloured according to whether they are within AA positions associated with differences in susceptibility to CTLs (pink) or susceptibility to NAbs (blue). The purple bar represents a probable CTL escape mutation, while the cyan bars represent selected variants within an epitope position targeted by the neutralising antibody HGF24. Where a codon is implicated in multiple responses, for clarity they are coloured in order of preference NAb, HGF24, CTL (see S2 Table).

### Distribution of selected variants among individuals

While the raw output of our inference method contains false positive inferences of selection, we can use statistical analysis to identify specific sites in the genome containing variants under selection. A unique aspect of this dataset is the large number of untreated individuals included in the study. Where alleles at the same locus were inferred to be under selection in multiple individuals, a statistical approach was used to infer loci at which, under conservative assumptions, at least one of the inferred variants is genuinely under selection. Taking into account different patterns for nonsynonymous and synonymous mutations, we estimated per-site false-positive rates for nonsynonymous and synonymous mutations, which were subsequently used to identify with statistical confidence sites that were under selection in at least one individual in our dataset. Taking all 34 individuals into consideration, we calculated that in p24 we could be confident that a site is under selection in at least one individual if mutations were inferred to be under selection in at least five individuals, and/or if nonsynonymous mutations were inferred to be under selection in two or more individuals, while in gp41 we could be confident if at least five mutations and/or three nonsynonymous mutations were inferred to be under selection. Applying these criteria, we identified 11 specific nucleotide sites, representing 10 amino acid (AA) positions, under selection in p24, and likewise we identified 11 such sites, representing 8 AA positions, in gp41. All but four mutations at these AA positions represented nonsynonymous changes (see Table 1). Occasionally two different codons at the same AA position were found to be subject to selection in a single individual, but in general repeated inferences of selection at an AA position occurred in distinct individuals (S1 Table).

**Table 1:** Summary of amino acid positions containing sites under selection.

| Region | Amino Acid Position<sup>a</sup> | Sensitivity<sup>b</sup> | Nonsynonymous<sup>c</sup> |  |  | Synonymous |
| --- | --- | --- | --- | --- | --- | --- |
|  |  |  | Reversion | Escape | Neither |  |
| p24 | 215[35–38] | CTL | 0 | 2 | 1 | 2 |
| p24 | 219[38–41] | CTL | 1 | 0 | 1 | 0 |
| p24 | 223[38–41] | CTL | 2 | 4 | 2 | 0 |
| p24 | 228[40,41] | CTL | 1 | 1 | 0 | 0 |
| p24 | 242[41–46] | CTL | 2 | 0 | 1 | 0 |
| p24 | 252[37,41] | CTL | 1 | 2 | 0 | 0 |
| p24 | 286 |  | 1 | 1 | 0 | 0 |
| p24 | 302[47] | CTL | 2 | 0 | 0 | 0 |
| p24 | 310[48–51] | CTL | 1 | 1 | 0 | 0 |
| p24 | 312[37,47,49,52,53] | CTL | 3 | 2 | 0 | 0 |
| gp41 | 620[54–58] | NAb | 3 | 3 | 4 | 1 |
| gp41 | 624[54,55] | NAb | 3 | 2 | 4 | 0 |
| gp41 | 641[59] | HGF24 | 0 | 2 | 1 | 1 |
| gp41 | 644[59] | HGF24 | 2 | 0 | 1 | 0 |
| gp41 | 648[59] | HGF24 | 1 | 3 | 0 | 0 |
| gp41 | 655[60] | NAb | 1 | 3 | 0 | 0 |
| gp41 | 674[57,61–66] | NAb | 0 | 2 | 1 | 0 |
| gp41 | 677[57,63,64] | NAb | 1 | 3 | 0 | 0 |

<sup>a</sup>Amino Acid (AA) positions in *gag* (p24) or *env* (gp41) relative to the HXB2 reference genome
<sup>b</sup>Sensitivity to CTLs or NABs using the Los Alamos HIV database (http://www.hiv.lanl.gov; see Methods for further details). AA position 228 in *gag* is associated with compensatory mutations,
rather than affecting sensitivity to CTLs directly. Position 286 in *gag* occurs in an epitope position targeted by a common HLA allele in the Ugandan population and so is likely associated with CTL escape. Positions 644, 648 and 655 were not associated with sensitivity to NABs, but are in an epitope region recognised by NAb HGF25.
°A nonsynonymous change was classed as a reversion if the nucleotide changed towards the subtype-specific population-level consensus, as an escape if the nucleotide changed away from the subtype-specific population-level consensus, and as neither of these if there was a nucleotide change which remained different to the subtype-specific population-level consensus.

Using the Los Alamos HIV database (http://www.hiv.lanl.gov), we determined for each of these AA positions whether they have previously been associated with changes in CTL susceptibility or NAbs (see Methods, Fig 2, S2 Table). In p24, nine of the 10 AA positions identified have previously been associated with changes in CTL susceptibility and/or compensatory mutations. The remaining codon (residue 286 in HXB2 *gag*) lies within CTL epitopes recognised by human leukocyte antigen (HLA) alleles that are relatively common in the Ugandan population (A1101 and B27[67]) and therefore mutations at this site probably affect sensitivity to CTLs, or possibly compensatory mutations. In gp41, of the eight AA positions identified as being under selection, five (residues 620, 624, 655, 674 and 677 of *env*) have previously been associated with NAbs, including broadly neutralising antibodies (bNAbs). These are associated with the gp120/g41 interface and the membrane-proximal external region (MPER). The other three codons (residues 641, 644 and 648 of *env*) all lie within an epitope region that is targeted by the monoclonal antibody HGF24, which has been shown to neutralise some viruses isolated from the African continent[59]. Although we cannot rule out other sources of selection in *env*, particularly due to CTLs, given the position of these codons on the genome, we believe selection due to humoral immune pressure is more likely. As such, all of the sites we identified as under selection in p24 probably affect susceptibility to CTLs or associated compensatory mutations, whereas all of the sites under selection in gp41 probably affect susceptibility to NAbs or associated compensatory mutations.

**Fig 2:**
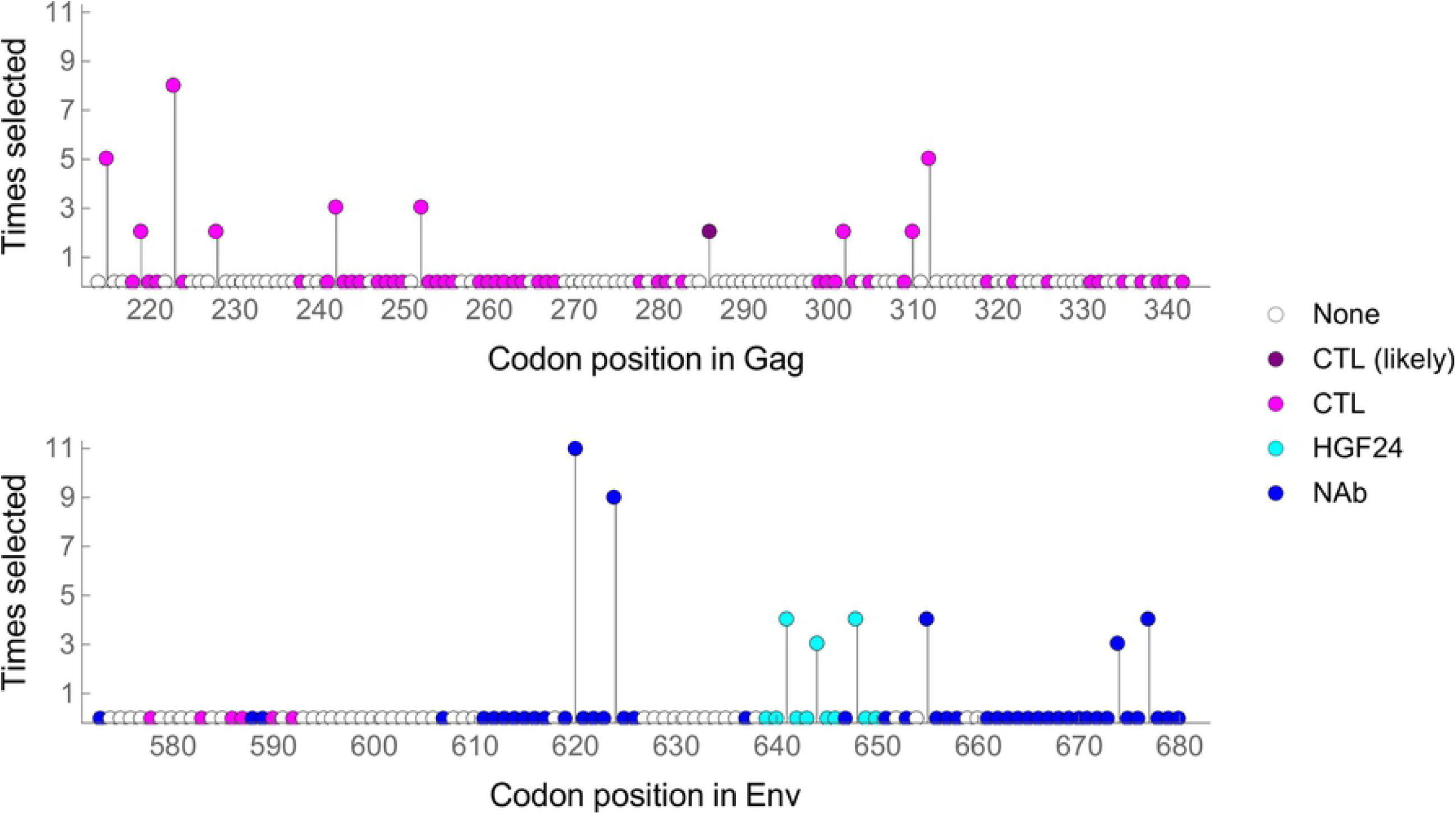
Amino Acid (AA) positions containing nucleotide sites that with confidence were inferred to be under selection in at least one individual. Codon positions are in relation to the HXB2 reference sequence. and the vertical y-axis gives the number of times that codon was inferred to be under selection across 34 individuals. Occasionally the same AA position was inferred to be under selection twice in the same individual. Pink: AA positions associated with changes in susceptibility to CTLs or compensatory mutations; Purple: AA probably affects susceptibility to CTLs; Blue: AA positions associated with susceptibility to NAbs; Cyan: AA positions in an epitope position targeted by the neutralising antibody HGF24. Where an AA position is implicated in multiple responses, for clarity they are coloured in order of preference NAb, HGF24, CTL.

### Direction of selection

Since mutations that enable the virus to evade host immune responses are often costly in terms of viral replication[68,69], sites harbouring escape mutations are likely to evolve towards the population-level consensus when transmitted to a new host since the selective pressure of a specific immune response is removed, but costs associated with the mutation remain [2,32,70–73]. This hypothesis is supported by the observation that allelic substitutions during the course of untreated infection occur towards population consensus much more frequently than expected by chance [2,32]. If analysis is restricted to sites where substitutions are observed, this bias is confined mainly to nonsynonymous substitutions[32], which is expected if immune escape mutations are generally nonsynonymous.

For the nucleotide sites that were confidently inferred to be under selection, we classified selected variants as escapes if they resulted in an AA change away from the subtype-specific population-level consensus in Uganda during the period the individuals were being sampled (see methods for full details). Conversely, inferred selected variants were classified as reversions if they resulted in change towards the subtype-specific population-level consensus. Of the 32 selected nonsynonymous mutations identified in p24, 13 were classified as escapes, 14 as reversions, and five as neither. The similar number of escapes and reversions observed is expected if CTL immune escape mutations are costly in non-HLA matched hosts, with escape in one individual followed by reversion in the next. Of the 40 nonsynonymous mutations identified in gp41, 18 were classified as escapes, 11 as reversions, and 11 as neither. The fact that over a quarter of AA changes are towards population consensus suggests that in many cases antibody escape mutations are deleterious in hosts without a matching antibody, but the stereotypical pattern of adaptation in one individual followed by reversion in another at CTL immune epitopes is likely more complex for antibody-escape evolution, where escape mutations are sometimes, but not always costly in the absence of antibody response [74,75].

We also determined the direction of change for all variants inferred to be under selection, which will likely include variants under direct selection and variants changing in frequency due to hitch-hiking, and whether they represented synonymous or nonsynonymous changes (S3 Fig). We found a clear correlation between the number of times that selection was inferred at an AA position and the proportion of selected variants that were nonsynonymous (linear regression, p24 p=0.009, r^2^=0.85; gp41 p=0.007, r^2^=0.67). For gp41, a correlation was also found between repeated selection at an AA position and a pattern of evolution towards the population consensus (p=0.011, r^2^=0.63); around half of inferred selection events were towards population consensus at the AA position most frequently inferred to be under selection. Although for p24 the linear regression did not reveal a significant trend (p=0.400, r^2^=0.18), there is a distinction between AA positions selected 2 or more times, which have a high probability of being towards subtype-specific population consensus (44%), and AA positions selected once, which have only a small probability of being towards the consensus (5%). Our interpretation is that AA positions represented once in our analysis disproportionately represent mutations increasing in frequency due to hitch-hiking, with these mutations tending to be synonymous and away from population level consensus. AA positions represented multiple times, on the other hand, are more likely to represent immune escapes and reversions, and therefore tend to be nonsynonymous but with only around half of mutations away from consensus.

### Comparing codon diversity at the within-host and population scale

Our data showed a strong relationship between within- and between-host sequence diversities, where the diversity of codons was measured at each AA position. Within-host diversity was measured approximately three years after seroconversion for each of the 34 individuals, and the mean calculated. Diversity at the population scale was calculated as the mean of the diversities for each of the subtypes A, D, and C, using virus sequences from a large number of individuals living in Uganda around the same time as the 34 individuals in our study (see Methods). Consistent with previous studies [2], we identified a strong relationship between measurements of sequence diversity calculated at the within-host and population scales (Fig 3), with all AA positions found to be highly diverse at the within-host level also highly diverse at the population level. Moreover, all but one of the AA positions containing nucleotide sites that we are confident are under selection are also diverse at the population level (the exception being residue 302 in *gag*). Given that most changes at these sites probably reflect escape from host immune responses, compensatory mutations, or reversions of these escapes in subsequent individuals, diversity at the population scale at these AA positions is likely maintained by the differing selection pressures faced by variants in different hosts due to different immunological backgrounds.

**Fig 3:**
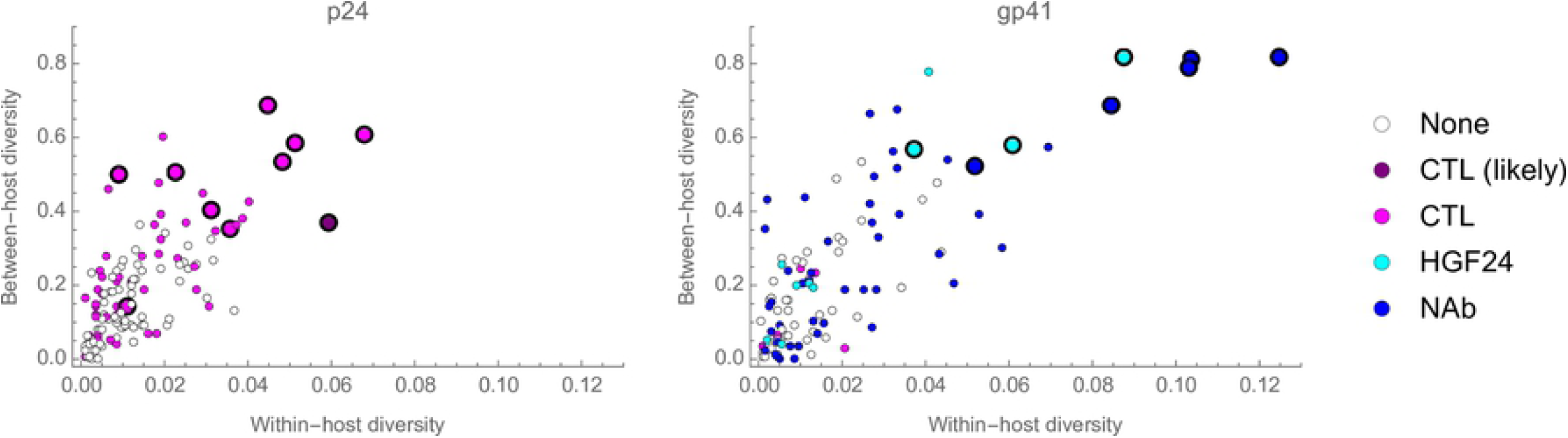
Within- and between-host codon diversity. For every AA position in our region of analysis we determined the within- and between-host codon diversity. Large markers denote AA positions in which we are confident selection is occurring in at least one individual. Markers are coloured if they are associated with changes in sensitivity to CTLs or compensatory mutations (pink) or NAbs (blue). In addition, AA postions in an epitope targeted by the neutralising antibody HGF24 are coloured in cyan. Where an AA position is implicated in multiple responses, for clarity they are coloured in order of preference NAb, HGF24, CTL.

Not all AA positions found to be highly diverse at the population level were detected as being under selection, with a high degree of confidence, at the within-host scale. This observation could arise if codons at these positions are under selection in some individuals in the population, but we failed to observe it with confidence, either because we sampled too few individuals, selection was too weak to be detected, and/or selection drove fixation events before the first sampling time point. In addition, at unconstrained sites experiencing little or no selection, diversity might gradually accumulate at the population scale due to drift, exacerbated by small bottleneck sizes at transmission. This could explain the high levels of population diversity at positions 235 in *gag* and 609 in *env*; almost all of the diversity observed at these codon positions is due to the presence of synonymous variants.

## Discussion

We developed a novel inference framework to infer the extent of selection acting upon variants which drive the evolution of within-host HIV-1 populations, considering data from the p24 region of *gag*, and gp41 of *env*, from 34 longitudinally sampled untreated individuals. A frequent assumption is that beneficial mutations will rapidly spread within individuals once they occur[19]. For example, it is well established that CTL-escape mutations accumulate and spread rapidly during acute infection [20–26], albeit that the rate of allele fixation decreases considerably during chronic infection[23,26]. However, estimating the extent and strength of positive selection during infection more generally is challenging due to genetic linkage among variant alleles, which makes differentiating between selected variants and variants that are increasing in frequency due to linkage with a selected variant (hitch-hiking) difficult. Our *de novo* approach incorporates genetic linkage and recombination. Furthermore, it is generally assumed that during untreated infection selected variants are associated with immune escape; our approach is agnostic with regards to phenotypic data, potentially allowing any polymorphic site in the genome to be identified as under positive selection.

Our results indicate a pattern of weak and slow selective sweeps characterising evolution during chronic HIV-1 infection, with stronger faster selective sweeps being relatively rare. We note that where ‘weak’ selection was inferred, this was still on a scale outweighing the effects of genetic drift. Studies of the effective population size of HIV-1 have indicated a value in excess of 10^5^ [76]; given such a value, selection of the order of 5% per generation is comfortably within a realm whereby the influence of selection dominates that of genetic drift[77]. An important caveat is that the first sampling time point for each individual in our analysis is estimated to be between 150 and 425 days since seroconversion, and therefore we will not detect variants that were under strong selection and rapidly reached fixation before the first sampling time point. Furthermore, the magnitude of the most strongly selected variants could not always be quantified; where fixation occurs entirely in the interval between two consecutive time points, no upper bound on the magnitude of selection could be fixed.

A unique aspect of our study is the large number of individuals for which we have data. Comparisons among individuals revealed AA positions which were inferred to be under selection in multiple (up to ten) individuals, with selected changes at these sites likely reflecting the gain or loss of CTL-escape mutations in p24, antibody-escape mutations in gp41, and escape-related compensatory mutations. Moreover, these sites were also found to be highly diverse at the population level. This is consistent with a pattern where a minority of codons are repeatedly under selection in multiple individuals, likely representing adaptation to the immunological background, which revert upon transmission to subsequent individuals; a pattern which has been referred to as “adapt and revert”[78]. Although we do not know the HLA-type of the infected individuals in our study, the number of putative CTL escapes and reversions is consistent with the frequency of different HLA alleles in the Ugandan population.

Perhaps less expected in our analysis was the identification of AA positions that are associated with neutralising antibodies and which were found to be under selection in a large number of individuals; for one site selection was inferred in nearly a third of individuals, with another inferred in a quarter of individuals. The implication is that the same epitopes are frequently targeted by antibodies in different individuals, and with similar means of viral escape. Moreover, since around a quarter of changes at these sites are towards the subtype-specific population level consensus, many may well represent the reversion of costly antibody-escape mutations from previous individuals, supporting the observation that some but not all antibody-escape mutations are costly [69,74,79–83]. These patterns can help explain why resistance to antibodies has increased over the course of the epidemic [84–88], but also highlights that viral evolution at the population level in response to bNAb-based interventions is likely to be complex, involving evolutionary responses to both naturally and therapeutically induced immune responses.

Even though our framework explicitly accounts for linkage disequilibrium between observed variants, it is still vulnerable to false positive inferences of selection due to linkage disequilibrium with unobserved variants flanking the genetic regions we analysed. Although simulated data suggested that the overall distribution of fitness effects was robust to this vulnerability, our study should serve as a cautionary note; where multiple alleles evolve in linkage disequilibrium, care is needed in identifying selection with any particular allele. The large number of individuals included in our study enabled us to partly circumvent this problem by only assigning confidence that any particular nucleotide site is under selection if it is inferred to be under selection in multiple individuals. Indeed, evidence for the validity of our method is provided by the repeated observation of variants under weak selection across multiple individual infections, with these changes making biological sense under the “adapt and revert” hypothesis. Our results emphasise the role of immune escape in driving evolution during chronic infection, shaping patterns of diversity at the population level, and provides new insights that could be useful in the development of immune-based interventions, particularly in the context of viruses circulating in Africa.

## Methods

In order to evaluate selection within a host, we employed a likelihood-based inference framework to infer the most parsimonious explanation of the sequence data in terms of a model of selection acting for specific nucleotides in the viral population. Some of the mathematical aspects of this framework have previously been applied in studies of the within-host evolution of the influenza virus[33,89], albeit that the model used in this case is specific to HIV-1. Our model explicitly accounts for linkage disequilibrium between alleles and builds upon earlier approaches for inferring selection in cases where linkage is of importance for evolution [24,28,90,91].

### Sequencing data

For our evolutionary analysis we used previously generated deep-sequence data from 34 longitudinally sampled individuals participating in the Rakai Community Cohort study and co-enrolled in the Molecular Epidemiology Research (MER) seroconverter study. Targeted short-read deep-sequence data from the p24 region of gag (390 bp; HXB2 reference genome positions 1429-1816) and the gp41 region of env (324 bp; HXB2 7941-8264) had been sequenced using the 454 sequencing platform (Roche, Branford, CT). All individuals were untreated, with a first sampling time point around one year since seroconversion, and typically 3 or 4 subsequent time points spanning between two and seven years of infection (see Table 1 in Raghwani et al 2019). Aligned sequences can be found at https://github.com/katrinalythgoe/ RakaiHIV. Further details on the individuals and sequencing methods used have been given elsewhere[32,92].

### Calling of variants from sequence data

As a preliminary step in the evaluation of sequence data, an estimate was made of the extent of noise in the data. While short-read sequencing allows very high read depths to be achieved, the incomplete sampling and inaccurate sequencing of a viral population imply that the extent of information in a viral dataset is often less than that assumed by a binomial or multinomial model[93]. While sequencing noise may be complex in nature, one approach to accounting for noise within an analytical likelihood framework is to employ a Dirichlet multinomial likelihood function with parameters equal to a constant term, C, multiplied by the underlying genotype frequencies[33]; the constant defining a generalised overdispersion in the data relative to a multinomial model. In the case of within-host influenza evolution a conservative estimate of this parameter may be generated by fitting a model of neutral evolution to a subset of genotype frequencies which are believed not to change substantially during the course of infection[33]; the inferred variance of the model, accounting for the read depth of sampling, provides an estimate of the extent of noise in the data.

HIV-1 infection occurs over a substantially longer period than does influenza infection, requiring a new strategy for the estimation of noise; we here exploited effects caused by genetic hitch-hiking[94]. If two alleles appear uniquely upon a shared genetic background, they will initially share an identical allele frequency. Over time the allele frequencies will change in a very similar manner, differences arising over time as a result of recombination between distinct haplotypes. We used putatively hitch-hiking trajectories to derive an estimate of noise in the sequence dataset. Loci were identified within the data from each patient at which a minor allele frequency of at least 10% was observed for at least two points in time. In a given patient we denote the trajectory

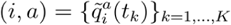

comprising the observed frequencies of allele *a* at locus *i* across all recorded times *t*_*k*_. For each pair of identified loci (*i, j)*, we found the alleles *a*, b\** in {A, C, G, T} minimising the statistic

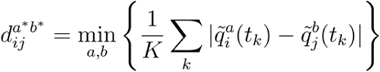

Initially, pairs of trajectories (i,a*) and (j,b*) were denoted as being ‘similar’ if

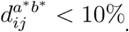

A set of further heuristic steps was then applied to refine these sets of trajectories. On timescales close to those over which the data for this study was measured, recombination in HIV-1 has been noted as being of importance over genetic distances greater than 100 nucleotides [2]; here a distance cutoff of less than or equal to 50 nucleotides was imposed between trajectories.

Next, pairs of variants were required to have similar frequencies at the first time of observation, requiring that

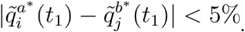

Further, so as to remove pairs of trajectories for which only one was polymorphic at a given time, it was required that the maximum ratio between minor allele frequencies did not exceed 10^3^ for the duration of the trajectories. Remaining sets of pairs identified in each patient were clustered into sets via an iterative process; beginning with an initial pair of trajectories *(i*_*1*_, *a*_*1*_) and *(i*_*2*_, *a*_*2*_), a trajectory *(i*_*m*_, *a*_*m*_) was added to the set if, following filtering, *(i*_*m*_, *a*_*m*_) was similar to a trajectory *(i*_*n*_, *a*_*n*_) already in the set. Sets of trajectories generated by the above process are shown in S4 Fig.

Having identified sets of trajectories, an inference process was used to evaluate the extent of noise in the data. Conservatively, the ‘true’ allele frequencies of each set were calculated as a simple mean of the observations, thereby assuming that all differences in frequencies result from ‘noise’ in the sequencing process. Given a set of trajectories *S* from a single viral population, where the allele frequency 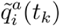 was calculated as 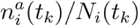, the frequency of observations reporting the given allele, the inferred frequency at time *t*_*k*_ was calculated as

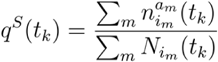

A Dirichlet multinomial model was then parameterised across all trajectory sets, finding the value of *C* satisfying

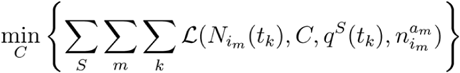

where

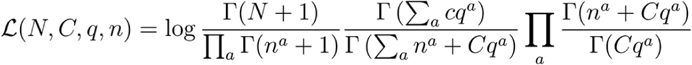

The derived value of *C* provides a proxy measurement of the extent of noise in the data and was used in further likelihood calculations; we inferred the value C = 100.359. We note that patterns of noise in genome sequence data may be substantially more complex than represented by our model; our likelihood, combined with the BIC model selection framework, provides a simple yet analytically tractable approach for the inference of selection parameters from real genome sequence data.

### Calling of variants from sequence data

Single-locus variants were identified in the data using the SAMFIRE software package[95]. Variants with a minimum allele frequency of at least 1% for at least one time point in the course of infection were identified. Considering frequency data from each variant allele, ‘potentially nonneutral’ loci [91] were identified, corresponding to positions in the genome at which significant changes in allele frequency were identified.

Variants were then filtered, identifying ‘potentially non-neutral’ loci [91] at which significant changes in allele frequency were observed over time. In this process, where q^1^(t) denotes the frequency of the variant allele at locus *i* at time *t*, deterministic models were fitted to the single-locus trajectories, using the equation

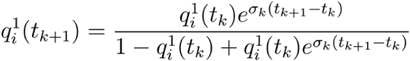

for neutral (σ_*k*_=0), constant (σ_*k*_=σ), and time dependent models of selection, retaining trajectories for which the constant or time-dependent models of selection outperformed the neutral model when evaluated using the Bayesian Information Criterion (BIC) to account for the increased complexity of the models including selection [96].

Targeted sequence data were processed into representative haplotypes, expressing each read in terms of the alleles present at the set of potentially non-neutral loci identified for that gene and individual. The evolution of the system was subsequently assessed in terms of the changes in frequencies of these haplotypes.

### Evolutionary model

Our evolutionary model considered the effect of mutation, selection, and recombination upon the population. Within the model the frequency of the haplotype *a* at generation *t*_*k*_ was specified by the frequency q_a_(t_k_), frequencies changing according to the three evolutionary processes. The model system was propagated within the space of observed haplotypes using a Wright-Fisher approach of discrete generations, with successive steps of mutation, recombination and selection.

Mutation was approximated as occurring between haplotypes that differ by a single nucleotide with rate μ per generation. Recombination was approximated as occurring in a pairwise manner between haplotypes with rate ρ per base per generation. That is, if a recombination event occurring between the loci indexed i and i+1, and involving the haplotypes a and b, were to produce the haplotype c, then in our model the new haplotype was produced at rate Δ_i,i+1_ρq_a_(t_k_)q_b_(t_k_) where Δ_i,i+1_ is the sequence distance between loci i and i+1.

A time-dependent model of selection acting upon haplotypes was applied, simulating the changing selection acting upon HIV during an infection. The time-dependent fitness *w*_*a*_ of a haplotype was modelled via a hierarchical model of single-locus terms

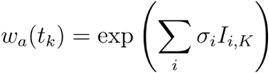

where the parameter σ_*i*_ denotes selection acting for or against all haplotypes with a variant locus at locus *i*, and the parameter *I*_*{i,K}*_ = 0 for all k ≤ K, and *I*_*{i,K}*_ = 1 for all *k* > *K*.

Selection then modifies the frequency of each haplotype according to the equation

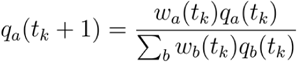

On the basis of previous studies[19,97–100], parameters for mutation and recombination were set at σ=3 x 10^−5^ per generation and σ=10^−5^ per base per generation, with a generation time of two days[101]. Parameters σ_*i*_ and *I*_*{i,K}*_, and initial frequencies *q*^*a*^*(t*_*0*_), were learnt from the data according to a hierarchical model framework. An initial, neutral model contained no parameters σ_*i*_. Next, single-locus selection models, with a single pair of selection parameters (σ_*i*_, *I*_*{i,K}*_), were considered; in each model the optimal values for these parameters were identified. By means of an iterative process, more complex models of selection were considered to the point of discovering a model for which neither adding an additional pair of selection parameters, or removing an existing pair of parameters, improved the model. As a conservative step, a model with an additional pair of selection parameters was required to significantly outperform a simpler model to be accepted, this being denoted by an improvement of 10 units of BIC upon the optimisation of model parameters.

The above model is an approximation in so far as the evolutionary process is evaluated only over the set of viral haplotypes which are observed in the sequencing data; the potential spread of the population into other haplotypes is not modelled. This approximation is necessary to make the calculation feasible; without such a constraint the exponentially large number of potential viral sequences would not allow model fitting to be performed.

### Estimates of uncertainty in selection coefficients

Error bars were calculated for each parameter σ_*i*_ inferred in the maximum likelihood calculation. Supposing the maximum log likelihood for a given inferred system to be equal to some value L, error bars were generated via a constrained exploration of the model space, in which a change in model parameters was accepted if the resulting likelihood was not greater than L-2, and for which changes to the parameter of concern, σ_*i*_, were constrained so that this parameter could only change in a specific direction; forcing this parameter to increase generated an estimate, after repeated iteration, for the upper error bar of this parameter, while forcing this parameter to decrease generated an estimate of the lower error bar of this parameter.

### Reporting selection coefficients

In our study we report the respective fitness advantage conferred by each beneficial mutation as a percentage per generation. This statistic, s, is calculated as

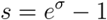

### Direction of Selection

For each of the selected mutations identified in our analysis, we determined whether evolution was towards or away from the subtype-specific population consensus. The subtype of each gene region for each individual and the population consensus for subtypes A, D and C in Uganda during a similar period to which the individuals were sampled was previously determined[32].

### Amino Acid position association with immune escape

Using the Los Alamos HIV database (http://www.hiv.lanl.gov), we determined for each of the AA positions in the p24 and gp41 regions that we analysed whether they have previously been associated with changes in CTL susceptibility or neutralising antibodies. For CTL susceptibility we used the Epitope Variant and Escape Mutation Database CTL variant search tool (https://www.hiv.lanl.gov/content/immunology/variants/variant_search.html?db=ctl). AA positions were marked as being associated with susceptibility to CTLs if susceptible and/or resistant variants were returned in the search tool. We excluded codons inferred to be susceptible to CTLs if the only evidence was high levels of diversity at the population level. For neutralising antibodies we used the Neutralizing Antibody Contacts and Features search tool (https://www.hiv.lanl.gov/components/sequence/HIV/featuredb/search/env_ab_search_pub.comp), marking codons as associated with susceptibility if variants were shown or predicted to affect neutralisation by antibodies, or binding to neutralising antibodies.

### Codon Diversity

We used the diversity statistic π to measure mean intra- and inter-host diversity at the codon level. Given an allele *a* with read depth *N*, where *n*_*i*_ of each of the codons *i ∈ {A,C,G,T}* have been observed, we define

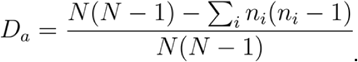

Then the diversity statistic π is defined by

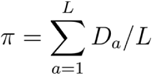

where the region of the genome under consideration is of length *L*.

This diversity statistic has been shown to be less prone than some other metrics to biases at the intra-host scale[102], and using codons (nucleotide triplet motifs) rather than amino acids or single nucleotides means our measure incorporates synonymous and nonsynonymous diversity whilst still enabling comparison with information on sensitivity to host immune responses, which is typically given at the amino acid level. To determine the mean intra-host diversity at a given codon position, we calculated π for each individual and then took the mean for all 34 individuals. To determine the mean inter-host diversity at a given codon position we calculated π for each subtype, and then took the mean for all three subtypes (A, C and D).

## Supporting Figures and Tables

S1 Fig: Codon positions containing nucleotide sites that are inferred to be under selection. This includes codons that are genuinely under selection and those that are increasing in frequency due to hitchhiking. Codon positions are in relation to the HXB2 reference sequence, and the y-axis gives the number of times that codon is inferred to be under selection across 34 individuals. Occasionally the same codon is inferred to be under selection twice in the same individual. Pink: codons associated with changes in susceptibility to CTLs; Purple: codon probably affects susceptibility to CTLs; Blue: codons associated with susceptibility to NAbs; Cyan: codons in an epitope position targeted by the neutralising antibody HGF24. Where a codon is implicated in multiple responses, for clarity they are coloured in order of preference NAb, HGF24, CTL.

S2 Fig: Distributions of input and inferred magnitudes of selection for simulated data in which the observed data described a fraction of the simulated region of the virus. Data are shown for variants at which the magnitude of selection could be inferred with confidence.

S3 Fig: Proportion of mutations inferred to be under selection that are towards population level consenus or are nonsynonymous. This includes codons that are genuinely under selection and those that are increasing in frequency due to hitchhiking. In all cases mutations are grouped according to the number of times the codon in which they appear is inferred to be under selection across the 34 individuals (x-axis). Top row: the number of codons in each group. Middle row: the proportion of mutations in each group that are towards population level consensus. Bottom row: the proportion of mutations in each group that are nonsynonymous.

S4 Fig: Sets of nucleotide trajectories that were identified as putatively hitchhiking. These trajectories were used to create a conservative estimate of the extent of noise in the sequencing data.

S5 Fig: True and inferred magnitudes and timings of selection for simulated data. Confidence intervals for the inferred selection coefficients are shown, calculated using the method described in the main text. The red dashed line indicates agreement between the true and inferred parameters.

S6 Fig: Observed (solid lines) and inferred (dashed lines) haplotype frequencies for simulated data in which all loci under selection were observed. In some cases the lines cannot be distinguished from one another.

S7 Fig: Observed (solid lines) and inferred (dashed lines) haplotype frequencies for simulated data in which only data from within a fraction of a simulated region was observed. In some cases the lines cannot be distinguished from one another.

S1 Table: Summary of results for all sites inferred to be under selection

S2 Table: Characteristics of all codons analysed: Sensitivity to host immunity, within- and between-host diversity, and the number of times the codon was inferred under selection

S3 Table: True and inferred selection parameters for a case in which sequence data describes all selected alleles within a system.

S4 Table: True and inferred selection parameters for a case in which sequence data partially describes the selected alleles within a system. Eleven out of 24 alleles were described in the data used for the inference calculation.

S1 Text: Calculations performed on simulated data, where targeted reads describe the complete evolution of the system, and where targeted reads do not describe the complete evolution of the system.

